# Endonuclease fingerprint indicates a synthetic origin of SARS-CoV-2

**DOI:** 10.1101/2022.10.18.512756

**Authors:** Valentin Bruttel, Alex Washburne, Antonius VanDongen

## Abstract

To prevent future pandemics, it is important that we understand whether SARS-CoV-2 spilled over directly from animals to people, or indirectly in a laboratory accident. The genome of SARS-COV-2 contains a peculiar pattern of unique restriction endonuclease recognition sites allowing efficient dis- and re-assembly of the viral genome characteristic of synthetic viruses. Here, we report the likelihood of observing such a pattern in coronaviruses with no history of bioengineering. We find that SARS-CoV-2 is an anomaly, more likely a product of synthetic genome assembly than natural evolution. The restriction map of SARS-CoV-2 is consistent with many previously reported synthetic coronavirus genomes, meets all the criteria required for an efficient reverse genetic system, differs from closest relatives by a significantly higher rate of synonymous mutations in these synthetic-looking recognitions sites, and has a synthetic fingerprint unlikely to have evolved from its close relatives. We report a high likelihood that SARS-CoV-2 may have originated as an infectious clone assembled *in vitro*.

**Lay Summary:** To construct synthetic variants of natural coronaviruses in the lab, researchers often use a method called *in vitro* genome assembly. This method utilizes special enzymes called restriction enzymes to generate DNA building blocks that then can be “stitched” together in the correct order of the viral genome. To make a virus in the lab, researchers usually engineer the viral genome to add and remove stitching sites, called restriction sites. The ways researchers modify these sites can serve as fingerprints of *in vitro* genome assembly.

We found that SARS-CoV has the restriction site fingerprint that is typical for synthetic viruses. The synthetic fingerprint of SARS-CoV-2 is anomalous in wild coronaviruses, and common in lab-assembled viruses. The type of mutations (synonymous or silent mutations) that differentiate the restriction sites in SARS-CoV-2 are characteristic of engineering, and the concentration of these silent mutations in the restriction sites is extremely unlikely to have arisen by random evolution. Both the restriction site fingerprint and the pattern of mutations generating them are extremely unlikely in wild coronaviruses and nearly universal in synthetic viruses. Our findings strongly suggest a synthetic origin of SARS-CoV2.

## Introduction

In just 2 years after SARS-CoV-2 emerged in late 2019, nearly 6 million people worldwide were confirmed to have died from COVID-19. Analyses of excess deaths estimate 18 million people lost their lives by December 2021 (Wang et al. 2022). Understanding the origin of SARS-CoV-2 can help managers prioritize policies and research to prevent future pandemics.

There are currently two hypotheses on the origin of SARS-CoV-2. The first hypothesis posits that SARS-CoV-2 has a natural origin and spilled over from animals to people at the Huanan seafood market (Pekar et al. 2022; Worobey et al. 2022). Research supporting a Huanan seafood market origin relies on analyses of early outbreak data suggesting the Huanan seafood market was an early epicenter of the COVID-19 pandemic. However, analyses of early case reports and phylodynamics are sensitive to assumptions about early case data. Such research assumes cases are ascertained at random, yet health authorities mounted extensive contact tracing and location tracing attempting to stop the early outbreak, and ties to the wet market were included as part of early case definitions (Washburne et al. 2022). Consequently, the wet market is believed to have been a site of transmission, it has not been conclusively shown to be the site of spillover.

The second hypothesis on the origin of SARS-CoV-2 posits that SARS-CoV-2 originated in a lab as a result of coronavirus (CoV) research. The lab origin hypothesis primarily notices that CoV research was carried out in Wuhan and that SARS-CoV-2 is unique among sarbecoviruses in having a Furin cleavage site (FCS) between the S1 and S2 subunits of the Spike protein. *In-vitro* studies have found the FCS is key to SARS-CoV-2 pathogenesis (Johnson et al. 2021).The FCS may explain why SARS-CoV-2 has caused a pandemic, while the estimated 66,000 sarbecovirus spillovers every year (Sánchez et al. 2022) do not. The FCS in SARS-CoV-2 is highly similar to one found in α-ENaC, a human epithelial Na channel gene (Anand et al. 2020; Harrison and Sachs 2022), which would be unusual for a sarbecovirus evolving in an animal host. However, the human-like ENaC is compatible with multiple explanations including lucky alignments, a non-human α-ENaC, acquisition of α-ENaC from post-spillover recombination, and more. More evidence is needed to discriminate between these two hypotheses and learn the origin of SARS-CoV-2.

Prior to the COVID-19 pandemic, many virological research projects examined how close naturally occurring CoVs are to causing a pandemic in humans. Researchers would explore the relationship between viral genotypes & human-infectivity phenotypes by a variety of experiments, including introducing small alterations generating Furin cleavage sites (Li et al. 2015) or experimenting with different receptor binding domains (RBDs) (Hu et al. 2017). Such experiments require making infectious clones, which requires assembling a full-length viral DNA genome *in vitro. In vitro* genome assembly (IVGA) has been used to create reverse genetic systems for many coronaviruses, such as transmissible gastroenteritis virus (Yount et al. 2000), MERS (Scobey et al. 2013), SARS (Yount et al. 2003), bat coronaviruses (Zeng et al. 2016), and more.

In this paper, we examine a common method for IVGA of RNA virus infectious clones. We document specific patterns in how researchers have historically modified viral genomes for IVGA. We find this specific pattern in SARS-CoV-2. We examine if the restriction map of SARS-CoV-2 meets all criteria for IVGA and estimate the probabilities of observed patterns in wild type CoVs as well as the odds of such patterns evolving from the close relatives of SARS-CoV-2.

## Results

The goal of this paper is to address the question whether SARS-CoV-2 originated from an animal-to-human spillover or from experiments performed in a laboratory. In the latter scenario, it is possible that evidence exists for manipulation of the viral genome by common laboratory techniques. SARS-CoV-2 is a large RNA virus. To create infectious versions of CoVs, the entire 30kb RNA genome is reconstructed in DNA by *in vitro* genome assembly (IVGA). IVGA has been used to create reverse genetic systems for modified and chimeric RNA viruses for more than 20 years (Yount et al, 2000). Most importantly, IVGA methods can leave genetic fingerprints, and we find those fingerprints in the genome of SARS-CoV-2.

### Methods and constraints for in vitro Genome Assembly

To make infectious clones from wild coronaviruses, one must synthesize a full-length DNA copy of the viral genome. Coronavirus genomes are ∼30Kb long. Making such a large DNA sequence requires assembly of smaller DNA fragments to create the larger, full-length viral genome. Assembly of larger DNA sequences from smaller segments can be accomplished efficiently using restriction enzymes that cut outside the enzymes’ recognition sequence, cleaving fragments of DNA and leaving unique 3-4 nucleotide overhangs with unique sticky ends permitting reliable reassembly of the fragments of DNA in the correct order (**Fig 1**). The full-length genome can be assembled in a bacterial artificial chromosome (BAC) or fragments stored separately in plasmids prior to assembly of a full-length cDNA (Almazán et al. 2014).

**Figure 1:**
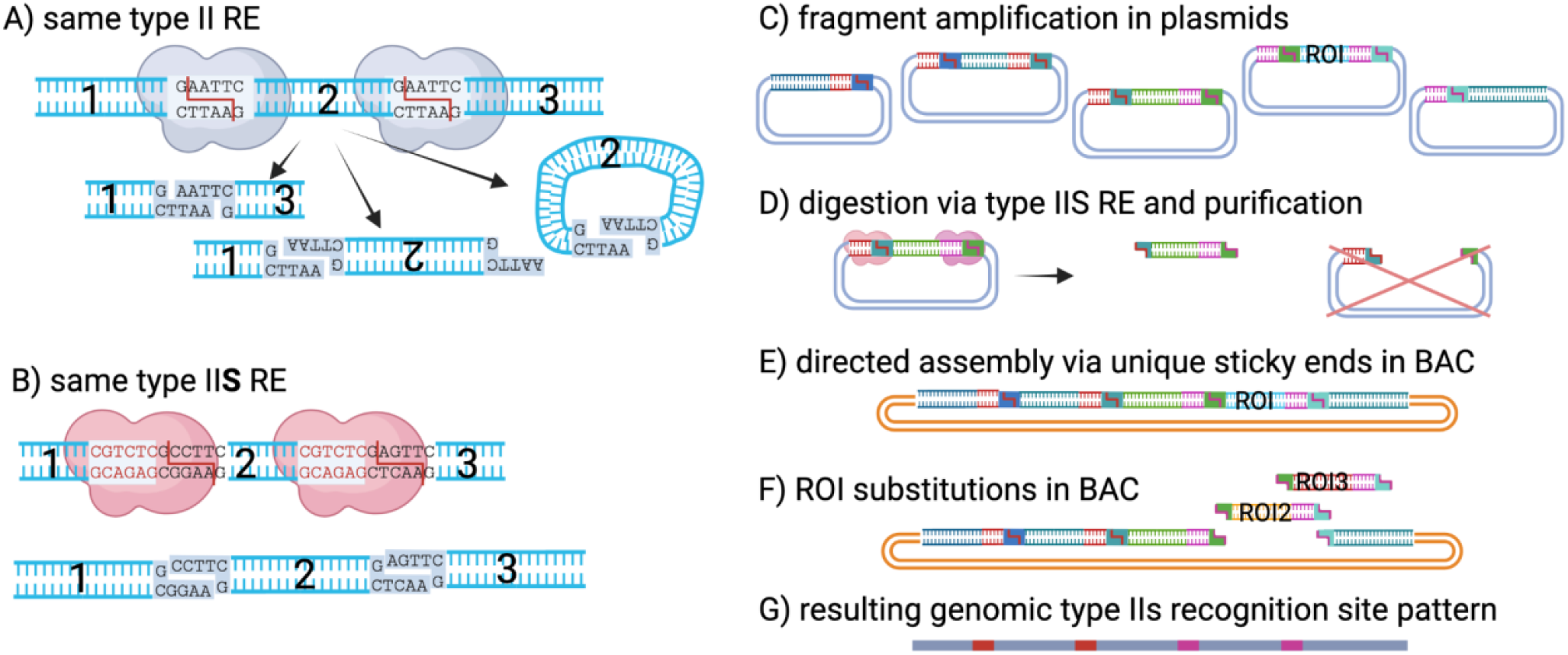
Synthetic RNA virus assembly. Directed assembly of ∼30kb CoV genomes requires several design considerations **A)** Several identical type II enzymes cannot be used for directed genome assembly as this leads to random fragment sequences, inverted fragments, and loops. Use of different type II enzymes that cut in their recognition sequence for every junction requires working with numerous buffers, running numerous reactions at different temperatures, and may require modifying numerous recognition sites in the genome. The use of fewer distinct enzymes is preferred. **B)** Endonucleases that cleave outside of their recognition sequence (type II shifted or type IIS) can produce distinct sticky ends allowing for directed assembly of complex viral genomes. **C)** For IVGA, individual fragments derived from PCR or DNA synthesis are first amplified in bacterial plasmids. **D)** Fragments are then cut out of the plasmids using type IIS endonucleases. **E)** Unique sticky ends at each section enable directed assembly in a full-length cDNA or bacterial artificial chromosome. **F)** Use of a different type IIS endonuclease with sites flanking a region of interest (ROI) allows for efficient substitutions of that region. **G)** This method does not alter viral proteins. However, it does leave a distinct pattern (fingerprint) of regularly spaced type IIS recognition sites of the endonucleases that were used for synthetic assembly.

IVGA has been used to create efficient reverse genetic systems to easily modify different segments of the viral genome and create chimeric coronaviruses to study the phenotypes of novel viral genotypes (Cockrell et al. 2017; Zhang et al. 2020; Messer et al. 2012; Donaldson et al. 2008; Donaldson et al. 2008). Making a reverse genetic system from a wild type CoV requires breaking the 30 kb coronaviral genome into 5-8 fragments, each typically shorter than 8kb (Almazán et al. 2006; Becker et al. 2008; Scobey et al. 2013; Zeng et al. 2016; Cockrell et al. 2017; Hu et al. 2017). To design a reverse genetic system, researchers often modify their synthetic DNA constructs from the wildtype viral genomes by introducing synonymous mutations that alter restriction enzyme recognition sites without significantly impacting the fitness of the resulting infectious clones.

Restriction sites are added and removed from the wildtype genome to partition the synthetic viral genome into several DNA segments that can each be mutated individually prior to assembling the full-length genome. While researchers could place restriction sites randomly throughout the genome, they instead tend to modify restriction maps in regular ways to accomplish research goals and meet the constraints of IVGA. Working with fewer fragments is easier, yet efficient fragment production requires all fragments not be too long. These protocol constraints result in regularly spaced restriction sites that minimize the number of sites and create a maximum fragment length that is shorter than expected by chance given the number of restriction sites. Re-assembled genomes typically lack scars, but the logistical constraints of infectious clone research results in regular spacing of restriction sites and a relatively small maximum length fragment, both of which become fingerprints of IVGA in the genomes of infectious clones.

The exact modifications of restriction sites are chosen to facilitate reseacrh goals. A 2017 publication introduced two BsaI sites into a bat CoV (WIV1) to enable efficient introductions of spike genes from other viruses (Hu et al 2017). The researchers used two distinct endonucleases for genome assembly, with two sites of one enzyme flanking a region of interest, enabling efficient manipulations of the flanking region without having to reassemble the entire viral backbone for each variant. A 2008 publication describes that restriction sites of a new SARS-like virus reverse genetic system were aligned with restriction sites of SARS reverse genetic system (Becker et al 2008). This could allow for efficient substitutions of segments between the two systems.

The following IVGA fingerprint can be observed in restriction site maps of synthetic viruses:

a. Introduction and/or deletion of unique endonucleases (BsaI, BsmBI, BglI).
b. Digestion with the chosen enzymes results in 5-8 fragments.
c. The largest fragment is less than 8 kb.
d. All sticky ends must be unique.
e. All recognition sites are created via synonymous mutations.
f. Two unique recognition sites may flank regions meant to be further manipulated.
g. Recognition sites may be aligned with other viruses to allow for segment substitutions.

### The genomic signature of IVGA

To investigate how these technical constraints and design considerations lead to distinct IVGA signatures, we first compute which random distributions of restriction sites can be expected in non-modified viruses. We do this by digesting a broad range of natural coronavirus genomes *in silico* with a comprehensive set of restriction enzymes to obtain a regular, “wild type distribution” of the maximum fragment length as a function of the number of fragments. The genomic signature of IVGA includes a specific type of outlier of the wild type distribution.

In 2013, researchers constructed a recombinant MERS coronavirus (Scobey et al. 2013). The wildtype virus had a few BglI sites at inconvenient locations, making it poorly amenable to efficient assembly. To construct an idealized MERS-CoV reverse genetic system for IVGA, the researchers removed the existing BglI sites and inserted 6 more evenly spaced BglI sites. All additions/removals were done via synonymous mutations, creating 7 fragments, the longest of which was 5721bp, or 19% the length of the 30kb MERS genome (**Fig 2A**). Under the wild type distribution, the average length of the longest segment for digestion producing 7 fragments was 40% of the genome. Researchers engineering MERS-CoV for IVGA sought evenly spaced type IIS restriction sites, leaving a fingerprint of longest-fragments that were unusually short compared to random wild type digestions.

**Figure 2:**
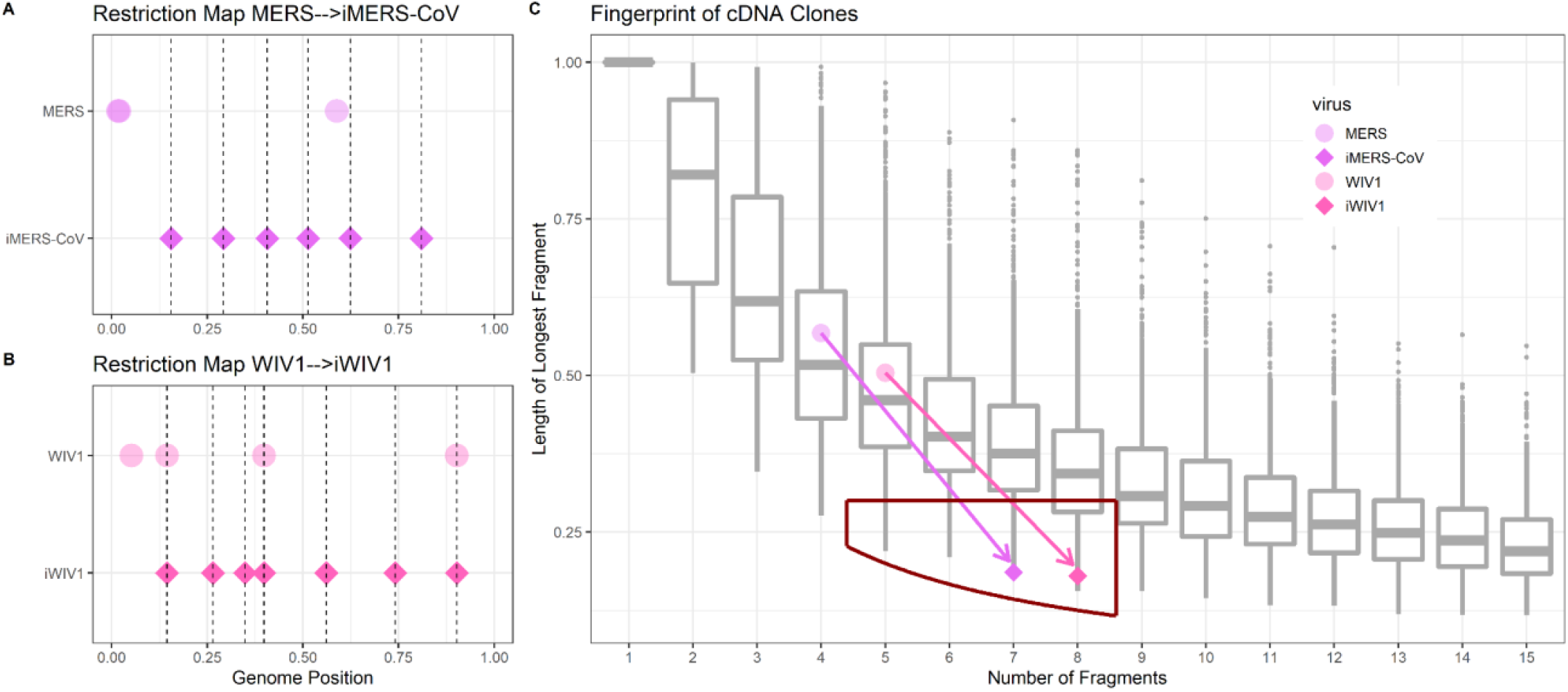
The restriction site fingerprint of *in vitro* genome assembly. **(A)** Compared to the wild-type genomes, a MERS virus engineered for IVGA, iMERS-CoV, has evenly spaced restriction sites, as does **(B)** a similarly engineered bat CoV, iWIV1. **(C)** *in vitro* assembled viruses deviate from the wild type distribution (gray boxplots) in identifiable ways. Due to research goals and laboratory logistical constraints, the longest fragments used to assemble cDNA clones are often significantly shorter than expected by the wild type distribution and the number of fragments remains low (5-8). To control for complex constraints on genomes, the wild type distribution of the longest-fragment length is estimated by digesting a wide range of non-engineered CoV genomes with a large set of endonucleases.

The same pattern can be seen in a modified SARS-like coronavirus. In 2016, researchers engineering a recombinant variant of the bat sarbecovirus WIV1 (iWIV1) utilized 3 pre-existing BglI sites, removed one pre-existing BglI site and introduced 4 new BglI sites, all through synonymous mutations (Zeng et al. 2016). iWIV1 was assembled from 8 fragments, and the maximum fragment length was 5451 bp (**Fig 2B**). Under the wild type distribution, the average longest-fragment length from a restriction digestion resulting in 8 fragments was 37% the length of the viral genome. Like iMERS-CoV, Infectious clones of iWIV1 had a longest-fragment from BglI digestion that was unusually short.

The effect of adding/removing type IIS sites for IVGA is shown in **Figure 2C**. The wild type distribution is used to estimate the likelihood of finding such anomalously short longest-fragment lengths in natural CoVs. Efficient reverse genetic systems used for IVGA prior to the emergence of SARS-CoV-2 have type IIS restriction maps with a narrow range of numbers of fragments and significantly shorter longest-fragment lengths than expected from the wild type distribution.

### The bioengineering utility of BsaI/BsmBI for SARS-CoVs

The most common restriction enzyme used for early IVGA approaches of CoVs was BglI (Yount et al. 2003; Scobey et al. 2013; Becker et al. 2008; Yount et al. 2000; Zeng et al. 2016). However, while MERS and SARS have several suitably located BglI sites in their genomes, the close relatives of SARS-CoV-2 have only one conserved BglI site that is inconveniently close to the beginning of the genomes (**Fig S1**). Also, BglI creates only 3nt overhangs, while type IIS endonucleases can produce 4nt overhangs that increase the odds of being unique and make for more reliable assembly of recombinant viruses. Two other commonly used type IIS endonucleases also used for IVGA are BsaI or the isoschizomers BsmBI/Esp3I (Donaldson et al. 2008; Rota et al. 2003). We only refer to BsmBI here (See **Table S1** for comments on type IIS enzymes). Sarbecoviruses contain a rich set of highly conserved BsaI/BsmBI restriction sites that can be used to create chimeric CoVs across a wide phylogenetic range of natural coronaviruses (**Fig 3B**). For bioengineers eager to study chimeric coronaviruses, BsaI/BsmBI would be an ideal combination to generate a flexible reverse genetics system from a natural SARS-CoV-2 related virus. To avoid losing power with multiple comparisons, we focus our analysis on the BsaI/BsmBI sites in SARS-CoV-2 and compare the BsaI/BsmBI map in SARS-CoV-2 to all other restriction maps of all other CoVs used in our analysis.

**Figure 3.**
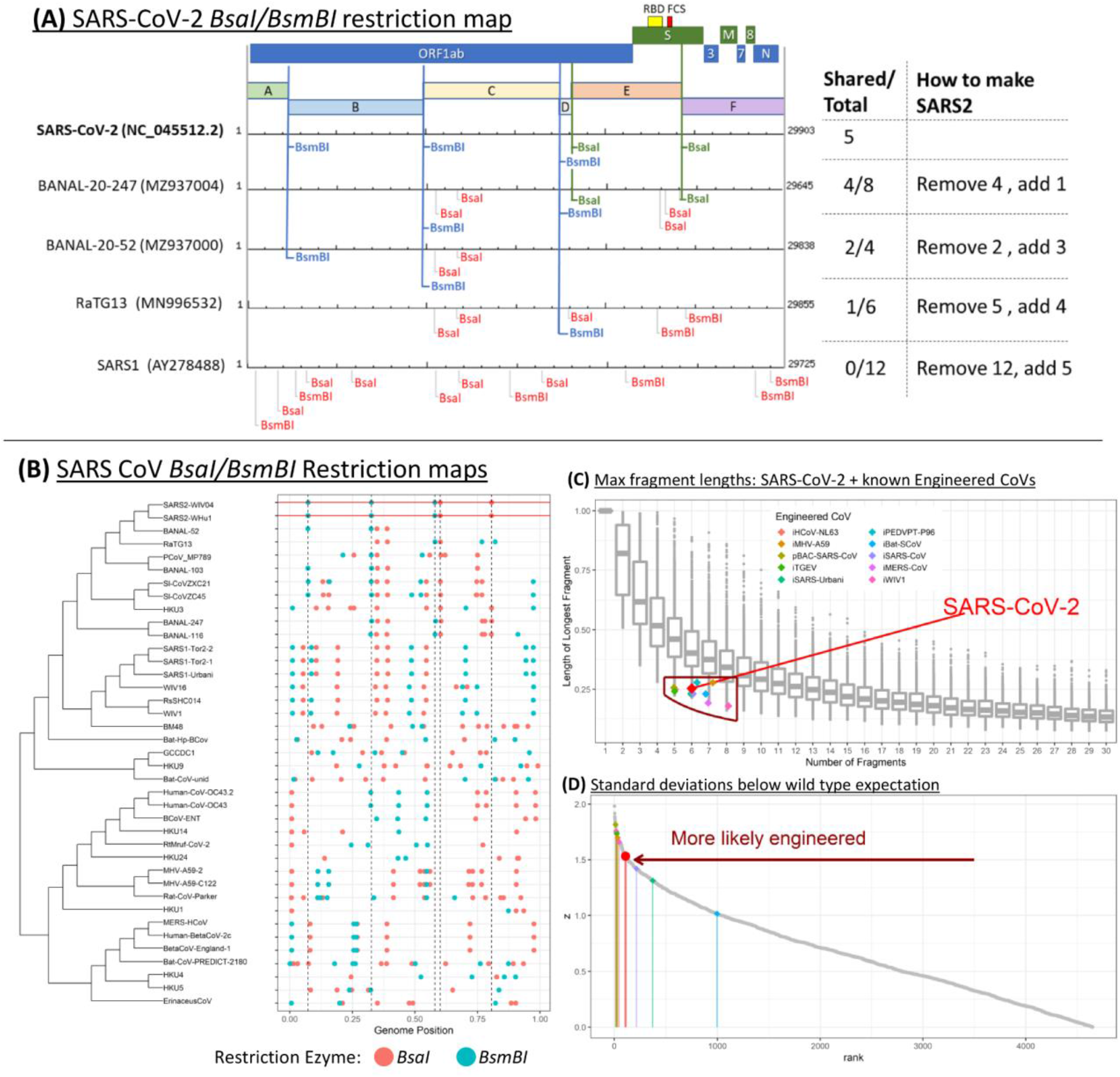
**(A)** How to make a SARS-CoV-2 BsaI/BsmBI restriction map from close relatives. **(B)** BsaI/BsmBI restriction map of the SARS coronaviruses. SARS-CoV-2 restriction sites are indicated with vertical dashed lines for comparison with other SARS COVs. Type IIS sites in identical positions in related viruses enable efficient substitutions of viral fragments to study chimeric CoVs. Having two BsaI sites flanking the S1 region and RBD enables efficient substitutions of receptor binding domains or introduction of an FCS at the S1/S2 junction. **(C)** Gray boxplots show the empirical null distribution of longest-fragment length from all CoVs and all restriction enzymes in our study. Colored dots show longest fragment lengths of known coronavirus reverse genetic systems, as well as SARS-CoV-2. SARS-CoV-2 and other CoV reverse genetic systems all fall underneath the null expectations as researchers seek to reduce longest-fragment lengths for IVGA. **(D)** A ranked plot of z-scores for all digestions creating 5-7 fragments, the idealized range for a CoV reverse genetic system. z-scores measure the standard deviations below the wild type expectation, correcting for the number of fragments. SARS-CoV-2 appears more likely to have been engineered for IVGA than several known CoV reverse genetic systems.

### SARS-CoV-2 BsaI/BsmBI fingerprint indicates probable in vitro origin

The SARS-CoV-2 genome contains 5 BsaI/BsmBI sites (**Fig 3A**). Its BsaI/BsmBI map contrasts with its close relatives by its even spacing and its absence of two highly conserved BsaI sites found in almost all other lineage B sarbecoviruses (**Fig 2 and 3B**). The SARS-CoV-2 BsaI/BsmBI restriction map would permit efficient modifications of the receptor binding domain, such as introduction of a Furin cleavage site at the S1/S2 junction, and allow substitutions of segments from related viruses which share these conserved recognition sites.

The longest-fragment length from BsaI/BsmBI digestion of SARS-CoV-2 is 7578 bp or 25% the length of its genome. Under the wild type distribution, the average longest-fragment length from a 6-fragment digestion is 43% the length of the viral genome (**Fig 3C**). The BsaI/BsmBI restriction map of SARS-CoV-2 is an outlier in the bottom 1% of longest fragment lengths of non-engineered CoVs, and it is consistent with observations from previously published coronavirus infectious clones (**Fig 3C**). Within all CoV restriction maps producing 5-7 fragments, the BsaI/BsmBI map of SARS-CoV-2 is more standard deviations below the wild type expectation than 3 of the 10 published type IIS-assembled CoV infectious clones (**Fig 3D**).

Limiting our wild type distribution to only type IIS enzymes with 6-7 nt recognition sequences and 3-4nt overhangs yields 1,491 CoV type IIS digestions fall within the ideal range of 5-7 fragments. Of the 1,491 restriction maps in the type IIS wild type distribution, SARS-CoV-2 is the more standard deviations below the mean than any non-engineered virus we found, suggesting under 0.07% chance of observing such an anomalous restriction map with such a high z-score in a non-engineered, wild type virus (**Fig S2**).

Random digestion and double digestion of CoVs with type IIS restriction enzymes with 6 nt recognition sites yields a median of 14 fragments and only 12.5% of these wild type digestions fell in the idealized 5-7 range. SARS-CoV-2 has 6 fragments upon double digestion. Its close relatives have 5 (BANAL 52) and 7 (RaTG13) fragments with distinct restriction sites (**Fig 3A**).

All sticky ends from type IIs digestions must be unique for assembly in IVGA cloning. All 5 of the 4nt overhangs from *BsaI/BsmBI* digestion of SARS-CoV-2 are unique and non-palindromic. All mutations modifying *BsaI/BsmBI* sites must be silent for ideal infectious clones. All 12 distinct mutations separating RaTG13 *BsaI/BsmBI* sites from SARS-CoV-2 are silent, and all 5 mutations between BANAL-52 and SARS-CoV-2 *BsaI/BsmBI* sites are silent. Between these two close relatives, 14 distinct silent mutations separate SARS-CoV-2 *BsaI/BsmBI* restriction sites from those of its close relatives. There are significantly higher rates of silent mutations within *BsaI/BsmBI* recognition sites for both RaTG13 (P=9×10^−8^; OR=8.9; 95% CIs: 4.2-17.3) and BANAL52 (P=0.004; OR=5.2; 95% CIs: 1.6-13.3) compared to the rates of silent mutations in the rest of the viral genomes.

### Mutation Analysis

100,000 random *in silico* mutants were generated for both RaTG13 and BANAl-20-52. The number of substitutions was equal to each genome’s nucleotide difference from SARS-CoV-2 and specific nucleotides substituted in were drawn in proportion to nucleotide frequencies across all 3 genomes. Mutants were digested *in silico*, the number of fragments & longest-fragment length extracted, and z-scores computed. Only 1.2% of RaTG13 mutants resulted in a *BsaI/BsmBI* restriction map with a larger z-score than SARS-CoV-2. BANAL52 is the closer relative to SARS-CoV-2 by over 200 nucleotides, yet only 0.1% of mutants yielded z-scores as great or greater than SARS-CoV-2. It’s unlikely such an idealized reverse genetic system would evolve by chance from the close relatives of SARS-CoV-2 (**Fig 4**).

**Figure 4:**
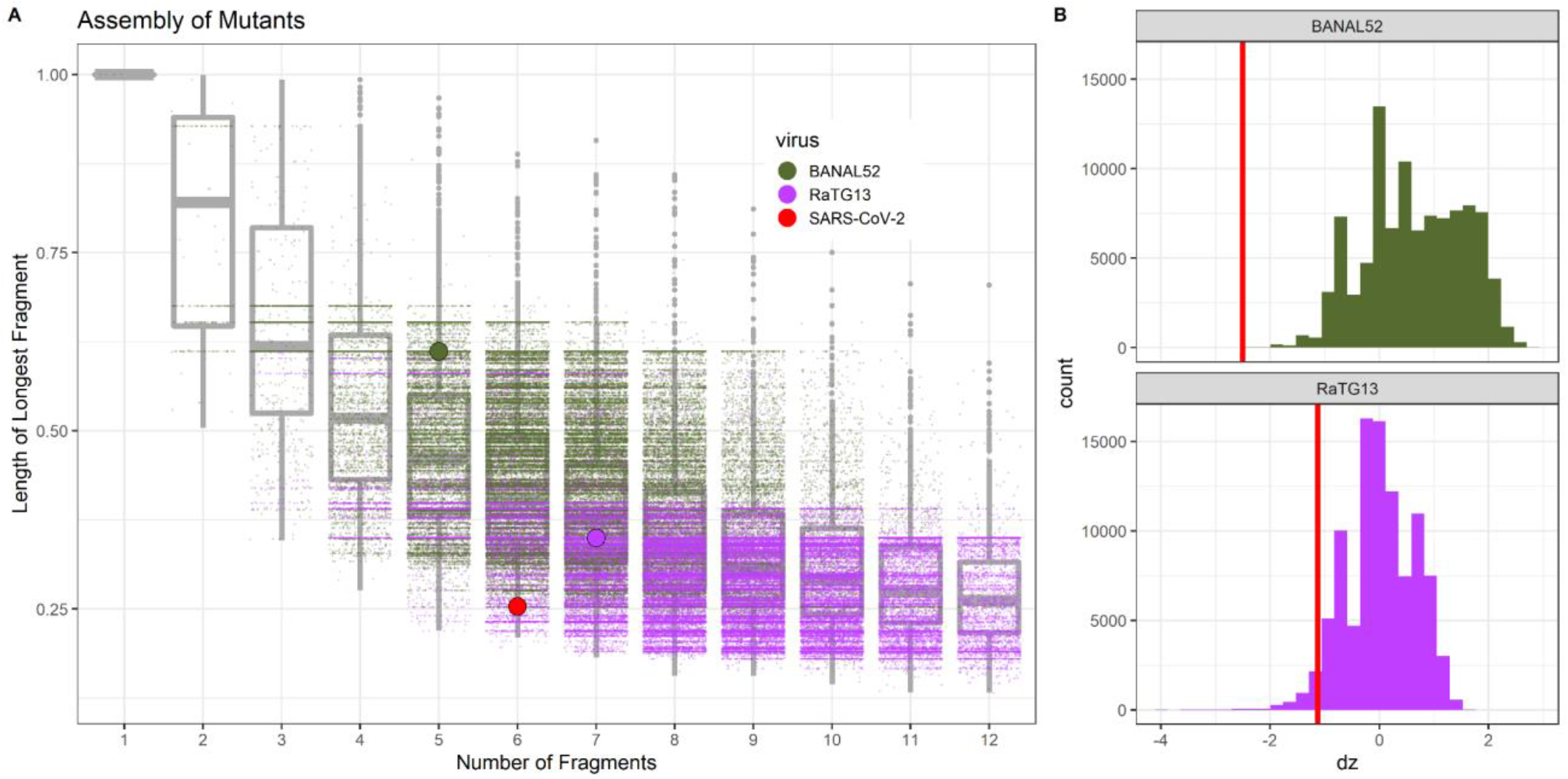
Mutation analyses of BANAL52 and RaTG13. **(A)** RaTG13 and BANAL-20-52 genomes were randomly mutated and digested by *BsaI/BsmBI in silico* to estimate the probability of natural mutations generating an infectious clone as good or better for IVGA than SARS-CoV-2. **(B)** We find a 1.2% chance of RaTG13 mutating to have a larger z-score than SARS-CoV-2, and a 0.1% chance of BANAL52 mutating to have a larger z-score than SARS-CoV-2.

Each alteration of BsaI/BsmBI sites between SARS-CoV-2 and BANAL-20-52 are caused by a single synonymous mutation in a wobble nucleotide, which is precisely how such alterations were made in the published studies described above. The combined odds of obtaining 5 wobble mutations by chance is likely very low (**Table S3**), although robust estimation of the odds requires considering a space of possible sites and careful examination of wobble mutation rates in the literature, so we leave this task to future research.

## Conclusion

The BsaI/BsmBI map of SARS-CoV-2 is anomalous for a wild coronavirus and consistent with an infectious clone designed as an efficient reverse genetics system. The research goals and laboratory logistics of infectious clone technology can leave a previously unreported fingerprint in the genomes of infectious clones. As a result of these constraints, published infectious clones have longest-fragment-lengths significantly shorter than those of natural CoVs digested by a range of restriction enzymes. The longest fragment in the BsaI/BsmBI restriction map of SARS-CoV-2 is in the bottom 1% of longest-fragments for all restriction maps analyzed. The longest fragment from BsaI/BsmBI digestion of SARS-CoV-2 is more standard deviations below the wild type expectation than any other non-engineered CoV digested by any IVGA-suitable type IIS enzyme in our analysis. When digested by BsaI/BsmBI, SARS-CoV-2 yields 6 fragments, falling within the idealized range for a reverse genetic system. The overhangs from BsaI/BsmBI digestion meet all the requirements for efficient and faithful lab assembly. All BsaI/BsmBI sites separating SARS-CoV-2 from its close relatives differ by exclusively synonymous mutations, with a significantly higher rate of synonymous mutations within BsaI/BsmBI sites than the rest of the viral genome. The BsaI/BsmBI restriction map of SARS-CoV-2 is anomalous among wild coronaviruses, consistent with a reverse genetics system, is unlikely to evolve from its closest relatives by chance. SARS-CoV-2 may have originated from a reverse genetics system.

The evidence we find is independent of other genomic evidence suggestive of a lab origin of SARS-CoV-2, such as the furin cleavage site (FCS) found in SARS-CoV-2 yet missing from all other known sarbecoviruses. However, the BsaI sites in SARS-COV-2 flank the S1 gene and S1/S2 junction, and a similar design has been used before for substitutions in this region. The restriction map alone also does not indicate a lab of origin.

Our theory that SARS-CoV-2 is a reverse genetics system can be tested. Databases of all CoVs collected and studied by relevant researchers may demonstrate that no progenitor to SARS-CoV-2 has existed in any known lab. Laboratory notebooks leading up to the November 2019 estimated start date of the COVID-19 pandemic may reveal no *BsaI*/*BsmBI* modifications of bat CoVs. A progenitor genome to SARS-CoV-2 found in the wild with the same or an intermediate *BsaI*/*BsmBI* restriction map may increase the likelihood of this anomalous restriction map evolving by chance.

Our analysis has several limitations. Our meta-analysis sought a representative set of engineered CoVs and searched for terms aimed to target specific literature using the specific method studied here. Expanding to other terms, literature, and the same methods applied to other viruses may improve our understanding of the fingerprint of IVGA. Additionally, our wild type distribution drew on a wide range of 70 non-SARS-COV-2 genomes and 214 restriction enzymes for sale at New England Biosciences, but restricting our analysis to just type IIS enzymes made SARS-CoV-2 an even larger outlier from the shifted wild type distribution. Additional CoV genomes and future research on null distributions of recognition sites may improve our understanding of the wild type distribution and lead to more robust quantification of the anomalous nature of the *BsaI/BsmBI* restriction map of SARS-CoV-2. We did not control for phylogenetic dependence among CoVs in our wild type distribution. Our mutation analysis considered a uniform rate of mutations across the genomes, whereas relative rates can increase or decrease the probability of reverse genetic system evolving from close relatives of SARS-CoV-2. Furthermore, we considered point mutations and not recombination events. While recombination events may explain portions of the SARS-CoV-2 *BsaI*/*BsmBI* restriction map, a formal analysis would require estimating the rate of recombination in CoVs, the size distribution of recombination events, the likelihood of the proposed recombination events in the evolutionary time between SARS-CoV-2 and its common ancestor with BANAL52, the distribution of possible ancestral states from recombination, and the combinatorial possibilities of recombination and mutation pathways that could generate SARS-CoV-2 under such an evolutionary model. These are important areas for future research.

Future research is also needed to better understand the evolution of restriction maps in CoVs. SARS-CoV-2 shares its first two BsmBI restriction sites with most other CoVs but not with RaTG13 nor the pangolin CoVs we found on NCBI. Meanwhile, the final three restriction sites of SARS-CoV-2 are not shared with most of the close relatives of SARS-CoV-2 but are found in distant CoVs like BANAL-20-247 and BANAL-20-113 (Temmam et al. 2022). Future research examining whole-genome evolution of restriction maps across a larger set of CoVs may produce more powerful tools to detect evolutionarily anomalous restriction maps.

Understanding the origin of SARS-CoV-2 can guide policies and research funding to prevent the next pandemic. The probable laboratory origin suggested by our findings motivates improvements in global biosafety. Given the advances in biotechnology and the low cost of producing infectious clones, there is an urgent need for transparency on coronavirus research occurring prior to COVID-19, and global coordination on biosafety to reduce the risks of unintentional laboratory escape of infectious clones.

## Materials & Methods

### Github Repository

Scripts used for analysis are stored at https://github.com/reptalex/SARS2_Reverse_Genetics. All analyses were conducted in R version 4.1.2. Scripts are numbered in the order of their implementation.

### Coronavirus Genomes

Coronavirus genomes for our phylogeny were obtained by using rentrez (Winter 2017). Spike gene ORFs were obtained by searching the NCBI Gene database for all Coronaviridae S-genes ORFs. Corresponding full genomes were pulled from the NCBI Genome database. An additional set of genomes was collected manually to ensure a balanced coverage of coronaviruses as important close relatives of SARS-CoV-2 were missing from our rentrez fetch. S-genes were extracted from the full genomes using the corresponding ORFs. Four of the Spike gene ORFs didn’t have corresponding genomes and were thus dropped. Four S-gene ORFs appear to be erroneous with nearly zero alignment with other CoVs, and were thus dropped, resulting in 72 Spike genes and corresponding full genomes for analysis. The resulting genomes, Spike gene sequences, and our alignment are all available on our Github repository.

### Engineered CoVs analysed in this study

Google Scholar searches were used to obtain a more complete and representative list of historical examples of infectious clones of coronaviruses. Three searches were conducted: {“coronavirus” “infectious clone” “type IIs”}, {“coronavirus” “infectious cDNA clone” “type IIs”}, and {“bat coronavirus” “infectious cDNA clone”}. Dates were limited from 2000-2019. All publicly available primary literature articles documenting novel infectious clones of coronaviruses were read and the following information was extracted: the virus, the wildtype accession number, number of fragments, maximum fragment length, and genome length. Literature reviews from our search were read and examples of coronavirus infectious cDNA or BAC clones mentioned were included. The resulting table of engineered CoVs used in our study is available on our Github repository.

### Phylogenetic Inference

Spike genes were translated with the R package Biostrings (Pages et al., n.d.) with input argument “solve” for fuzzy strings and then aligned on Mega X (Kumar et al. 2018) using ClustalW (Thompson, Gibson, and Higgins 2002). A maximum likelihood phylogeny was constructed with default settings (JTT substitution model, G+I rates, and NNI heuristic method for ML inference). Our phylogeny was constructed only for the purpose of enabling easy visualization of restriction maps of close relatives.

### Restriction map analysis

The R package DECIPHER (Wright, n.d.) was used to analyze restriction maps. DECIPHER comes with a set of 214 restriction enzymes for sale at New England Biolabs (referred to as NEB restriction enzymes). To obtain the null distribution of CoV restriction maps, we digest all 72 CoV genomes with each of the 214 restriction enzymes and 1,000 randomly drawn pairs of different restriction enzymes. The type IIs restriction maps in this set that were specifically amenable for

BAC cloning and in the NEB restriction enzyme set were: BbsI, BfuAI, BspQI, BsaI, BsmBI, and BglI. For type IIs digestions, we used all aforementioned type IIs enzymes and all distinct pairs of these enzymes.

The number of fragments *n* and maximum fragment lengths, *L*, expressed as a proportion of genome length, were extracted for analysis. For species *i* and restriction map *r* we obtained a maximum fragment length *L*_*i,r*_ resulting in *n* fragments, and z-scores were calculated

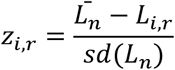

Where 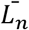 is the expectation *sd*(*L*_*n*_) is the standard deviation of maximum fragment lengths for all restriction maps across all species yielding *n* fragments.

### Mutation analysis

Whole-genome pairwise alignments between RaTG13 and SARS-CoV-2 and BANAL-20-52 and SARS-CoV-2 were implemented using MUSCLE (Edgar 2004). A number of random substitutions equal to the nucleotide difference between the genomes were simulated. Sites were selected at random and mutated to another base with probabilities proportional to the frequency of bases in the three CoV genomes. Mutant genomes were digested with BsaI/BsmBI and z-scores were extracted as described above.

Fisher’s exact test was used to assess if there was a higher rate of silent mutations within BsaI/BsmBI recognition sites compared to the rest of the viral genome. Odds ratios were computed as the ratio of silent mutations to all other nucleotides within BsaI/BsmBI sites of either genome in a pairwise alignment compared to the ratio of silent mutations outside BsaI/BsmBI sites to all other nucleotides. There are 12 silent mutations found in 9 distinct BsaI/BsmBI sites between RaTG13 and SARS-CoV-2, and 882 silent mutations outside of BsaI/BsmBI sites. There are 12 silent mutations found in 9 distinct BsaI/BsmBI sites between RaTG13 and SARS-CoV-2, and 882 silent mutations outside of BsaI/BsmBI sites. There are 5 silent mutations found in 7 distinct BsaI/BsmBI sites between BANAL52 and SARS-CoV-2, and 753 silent mutations outside BsaI/BsmBI sites.

## Acknowledgements

We thank Justin B. Kinney (Cold Spring Harbor Laboratory) for helpful discussions and for feedback on the manuscript. We thank many other unnamed colleagues for their feedback.

## Supplemental Information

**Figure S1:**
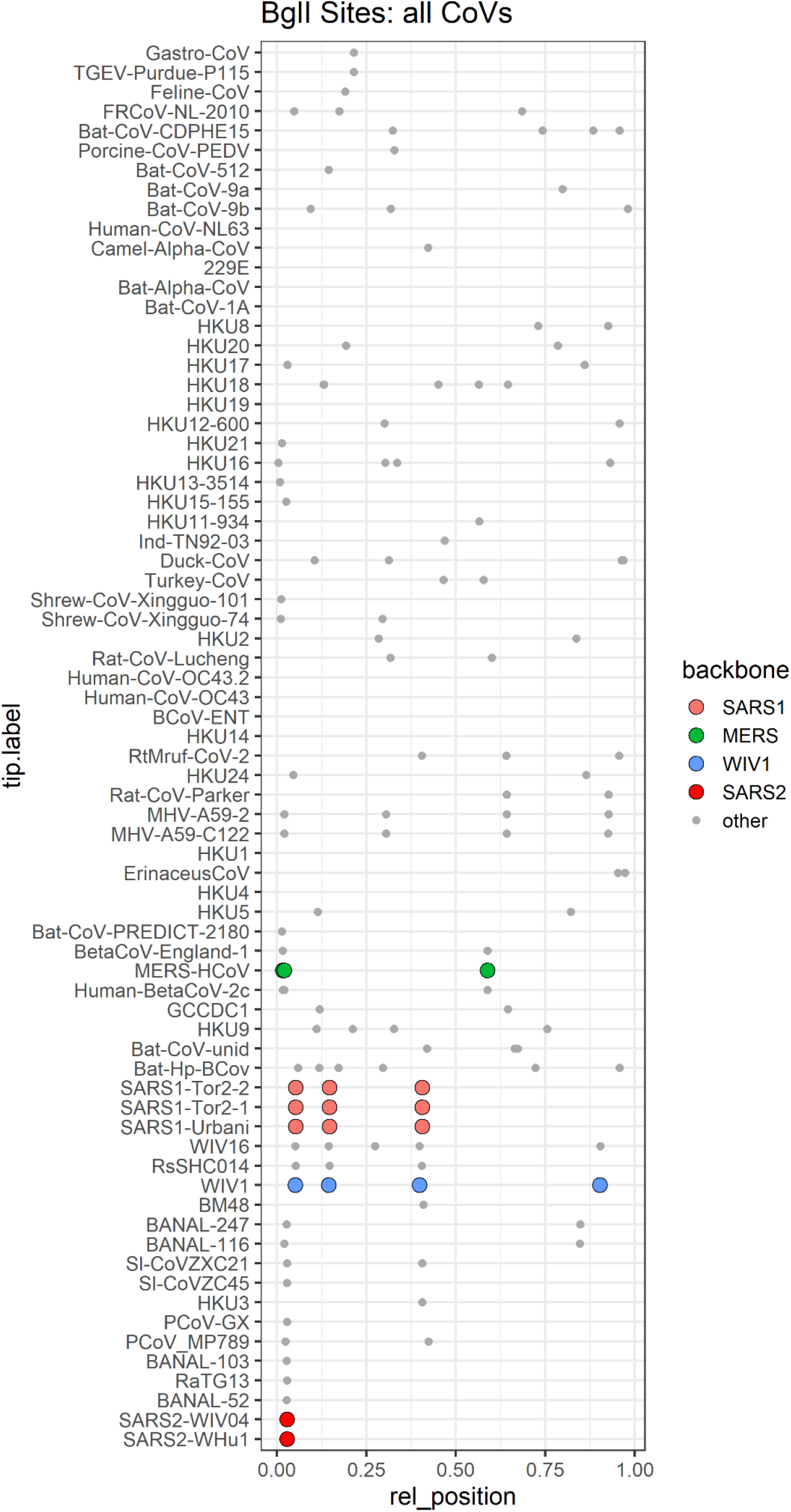
BglI sites for all CoVs, in order of their tip-labels on the broader CoV phylogeny. Colored dots indicate wild type viruses for which BglI was used to make recombinant viruses. In these viruses, BglI has several sites which are conserved in the crown group, allowing more natural construction of recombinant viruses based on conserved type IIs sites.

**Figure S2:**
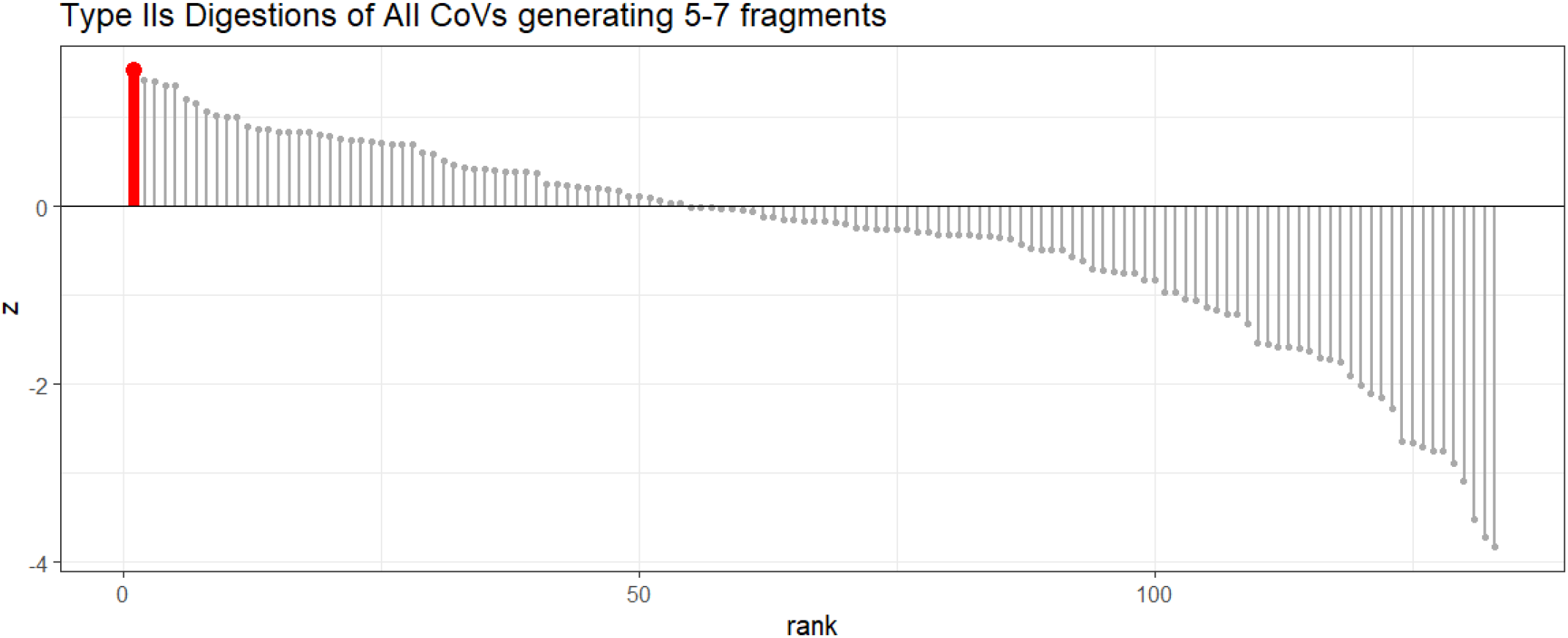
Z-scores for all CoV type IIs restriction maps falling within the idealized range (5-7 fragments) for an efficient reverse genetics system. Of 1065 combinations of CoVs and restriction enzyme digestions or double-digestions, the SARS-CoV-2 type IIs restriction map is the most anomalous. No other CoV we analyzed has a type IIs restriction map with the idealized number of fragments and a maximum fragment length more standard deviations below the mean than the BsaI/BsmBI site of SARS-CoV-2.

**Table S1:** disadvantages of alternative type IIS REs according to SnapGene ® software 2022

| Endonuclease | Potential disadvantage |
| --- | --- |
| BbsI | Instability over time |
| BfuAI and BspQI | require 50°C |
| BspQI and SapI | Produce only 3nt sticky ends |
| PaqCI | Has a 7nt recognition sequence, more difficult to introduce |
| Esp3I | Non, same recognition sequence as BsmBI, not distinguishable in retrospect |

Other type IIs RE can be used for IVGA, but have certain limitations, including instability over time, a higher required temperature for cleavage, only 3nt overhangs or a 7nt recognition sequence which can be more difficult to introduce without causing nonsynonymous mutations. Esp3I is an isoschizomere of BsmBI, a few more isoschizomeres are not listed here.

**Table S2:** Criteria for *in vitro* genome assembly and estimated probabilities in wild type CoVs.

| Criterion | Test | Probability/P-value | Notes |
| --- | --- | --- | --- |
| Anomalously Short max-fragment length | Z-score controlling for # of fragments | 1% (all REs)<br><0.07% (type IIS) | Sensitive to wild-type distribution |
| 5-7 fragments | % of type IIS digestions in range | 12.5% | Sensitive to wild-type distribution |
| All unique, non-palindromic overhangs that are not exclusively GC | Examine SARS-CoV-2 overhangs | 60% | Simulated random sets of 5 4mers with same base frequency as SARS-CoV-2 |
| All restriction site-modifying mutations are synonymous | Binomial test of synonymous mutations | $P = 0.08$ | Controlled for dN/dS heterogeneity across species, not across sites |
| High concentration of synonymous mutations per nt in RE sites | Fisher exact test | $P = 9 \times 10^{-8}$ (RaTG13)<br>$P = 0.004$ (BANAL52) | Did not control for dN/dS heterogeneity across genome |

**Table S3:**
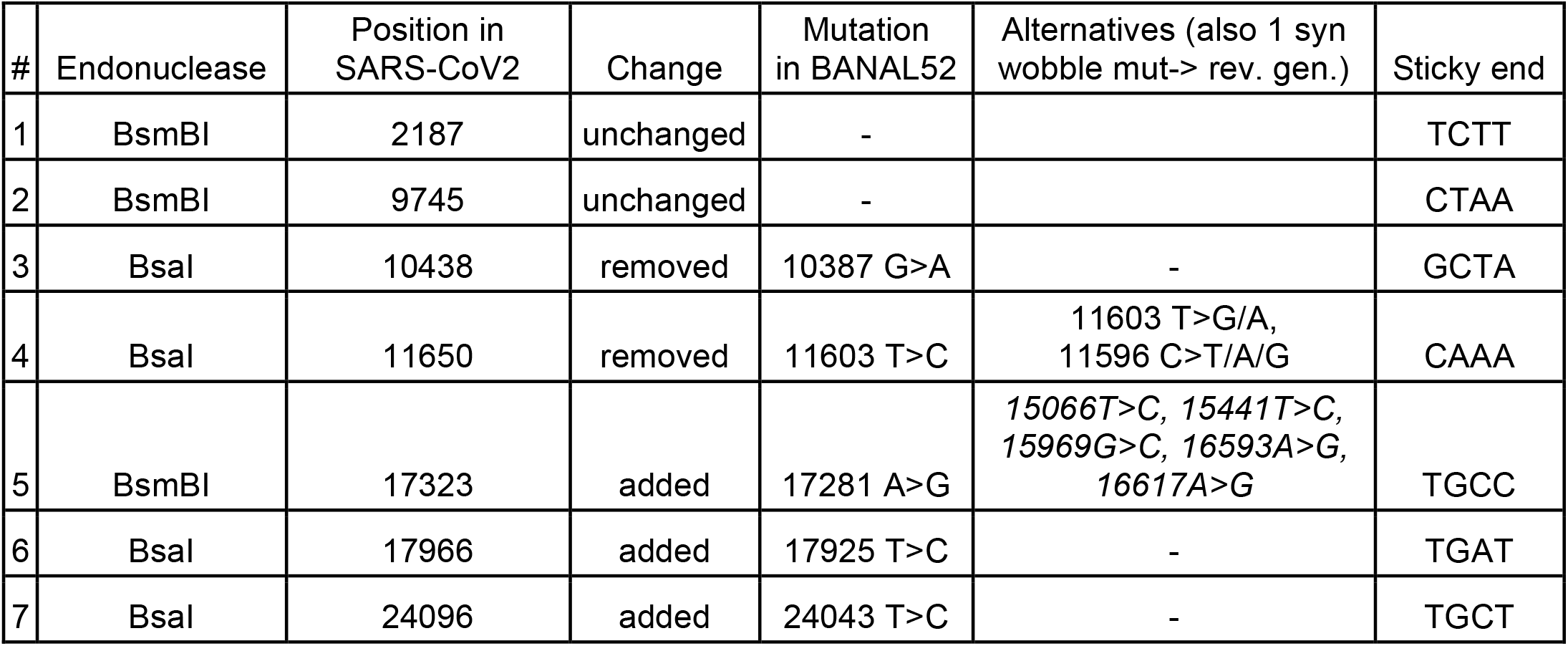
Overview BsmBI & BsaI sites in SARS-CoV-2 or Banal-20-52, cleavage position in SARS-CoV2, the mutation in BANAL-20-52 that would lead to the desired change, all alternative synonymous wobble mutations that would lead to an equally efficient reverse genetics system with only a single mutation, and the respective sticky ends.

